# Estimating the impact of city-wide *Aedes aegypti* population control: An observational study in Iquitos, Peru

**DOI:** 10.1101/265751

**Authors:** R.C. Reiner, S.T. Stoddard, G.M. Vazquez-Prokopec, H. Astete, T.A. Perkins, M. Sihuincha, J.D. Stancil, D.I. Smith, T.J. Kochel, E.S. Halsey, U. Kitron, A.C. Morrison, T.W. Scott

## Abstract

During the last 50 years, the geographic range of the mosquito *Aedes aegypti* has increased dramatically, in parallel with a sharp increase in the disease burden from the viruses it transmits, including Zika, chikungunya, and dengue. There is a growing consensus that vector control is essential to prevent *Aedes*-borne diseases, even as effective vaccines become available. What remains unclear is how effective vector control is across broad operational scales because the data and the analytical tools necessary to isolate the effect of vector-oriented interventions have not been available. We developed a statistical framework to model *Ae. aegypti* abundance over space and time and applied it to explore the impact of citywide vector control conducted by the Ministry of Health (MoH) in Iquitos, Peru, over a 12-year period. Citywide interventions involved multiple rounds of intradomicile insecticide space spray over large portions of urban Iquitos (up to 40% of all residences) in response to dengue outbreaks. Our model captured significant levels of spatial, temporal, and spatio-temporal variation in *Ae. aegypti* abundance within and between years and across the city. We estimated the shape of the relationship between the coverage of neighborhood-level vector control and reductions in female *Ae. aegypti* abundance; i.e., the dose-response curve. The dose-response curve, with its associated uncertainties, can be used to gauge the necessary spraying effort required to achieve a desired effect and is a critical tool currently absent from vector control programs. We found that with complete neighborhood coverage MoH intra-domicile space spray would decrease *Ae. aegypti* abundance on average by 67% in the treated neighborhood. Our framework can be directly translated to other interventions in other locations with geolocated mosquito abundance data. Results from our analysis can be used to inform future vector-control applications in *Ae. aegypti* endemic areas globally.

**Author Summary:** Despite the growing threat of arboviruses, there is a dearth of ‘best practices’ for the primary vector control tools used in the field. In the absence of cluster randomized control trials, evidence on the utility (or lack thereof) of vector control interventions must be gleaned from ongoing control programs. Motivated by 12 years of household-level Ae. aegypti abundance surveys and neighborhood-level space-spray campaign data from Iquitos, Peru, we developed a new framework to model mosquito abundance. In spite of significant spatial and temporal heterogeneity, we identified a statistically significant and practically important impact of the local Ministry of Health space-spray campaign, specifically, a reduction of mosquito abundance of 67% when coverage was optimal. Our framework can be directly applied to other locations with geolocated mosquito abundance data and our findings can be used to both optimize resources within Iquitos as well as inform future vector-control interventions in *Ae. aegypti* endemic areas globally.

## Introduction

*Aedes aegypti* is the primary vector for multiple arboviruses [1]. Controlling these pathogens is of paramount importance, especially with recent invasions into the Americas of chikungunya and Zika, outbreaks of yellow fever reviving concerns of urban yellow fever epidemics, and the suboptimal performance of the only licensed dengue vaccine [2]. Until better options arise, controlling transmission by reducing the vector population remains the primary tool available to combat the *Aedes*-borne disease burden.

A hemisphere-wide eradication campaign in the Americas in the 1950s and 1960s demonstrated that control of this mosquito can be a viable option for controlling disease from the viruses they transmit [3]. In the vacuum left by the cessation of that eradication program, which was likely exacerbated by urbanization and globalization of trade and human travel [4], *Ae. aegypti* recolonized significant portions of the Americas [5]. Current *Aedes*-transmitted disease control strategies lack the scale and support of the multi-country eradication effort. They are often developed and administered on a provincial or city-by-city basis and frequently ineffectively applied [6]. There is a notable lack of “best practices” to be followed [7] and an absence of rigorously collected, quantitative entomological data that can be used to inform vector control strategies [8]. Furthermore, resistance to various insecticides, particularly pyrethroids, further hinders the effectiveness of existing interventions [9]. Controlled laboratory experiments can assess the efficacy of an exact dosage of insecticide on a mosquito, but it is unclear how that translates to the real-world effectiveness of the application of that insecticide in, for example, a citywide space-spray campaign. Quantifying the effectiveness of real world control programs on mosquito populations is necessary to assess a program’s utility (or lack thereof) and to provide useful evidence for enhancing future intervention programs.

There are three key factors that complicate the estimation of an intervention’s impact on *Ae. aegypti* population size. First, *Ae. aegypti* abundance is highly heterogeneous in space and time. *Aedes aegypti* population dynamics are strongly influenced by micro-climate [10,11], they have a limited flight range (∼100m [12]), and they exhibit fine-scale spatial clustering (30m in urban areas [13,14]). Small seasonal fluctuations in weather or available larval habitats can affect *Ae. aegypti* population dynamics from year to year, or even month to month [14]. Variation in observed populations, therefore, can be just as likely attributed to environmental fluctuations as a mosquito control program.

Second, to calculate a control program’s effect size, it is necessary to estimate what would have happened without an intervention; i.e., the *counterfactual*. In many locations, resources are limited and are directed towards control as opposed to surveillance. In the absence of substantial surveillance, it is extremely difficult to parse the impact of an intervention from natural variations in mosquito populations [6].

Third, accurately measuring *Ae. aegypti* abundance is difficult even under optimal circumstances [15]. *Ae. aegypti* is typically found in low abundances in and around human habitations, with sample sizes averaging sample of 1-5 adults per house [16]. Larval surveys do not capture variation or magnitude of adult abundance and miss cryptic habitats [17]. Most *Ae. aegypti* control programs do not have the personnel and procedures in place to estimate spatial and temporal variation in population abundance across cities, districts, or states, although there are some notable exceptions [18]. Large-scale, household-level measurements are required across multiple years using a consistent, reliable measurement method to characterize baseline spatio-temporal patterns in abundance, which is seldom done [19]. Overcoming measurement error represents a significant challenge, because *Ae. aegypti* is a low-abundance species, adults are difficult to collect, and there are no reliable entomological measures for risk of human arboviral infection and/or disease [20].

Iquitos, Peru, a city of approximately 370,000 and the largest urban center in the Peruvian Amazon, is endemic for *Ae. aegypti* and dengue viruses. Between 1999 and 2010, 176,352 geolocated, household-level *Ae. aegypti* abundance surveys were conducted using hand-held mosquito aspirators (Figure 1), resulting in the capture of 48,015 female *Ae. aegypti.* Local Ministry of Health data was available on spatiotemporally explicit mosquito control efforts. The goals of the current work were two-fold. First, we developed a framework to model *Ae. aegypti* populations at a fine scale through space and time. Second, we used that framework to explore the impact of a city-wide vector control program by the Ministry of Health in Iquitos.

**Figure 1:**
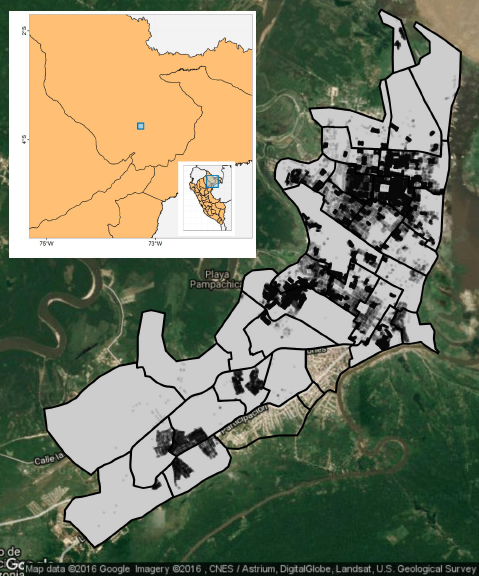
Collection locations and Ministry of Health Zones. Ministry of Health Zones are plotted with thick black outline. All but three had some level of mosquito collection (grey background), with each house-hold aspiration indicated by a black circle.

## Results

### Data overview

Summary statistics of mosquito data indicated a substantial effort to capture mosquitoes each month; more than 500 individual female *Ae. aegypti* mosquitoes were captured during most months (Figure 2a). The average number of captured female *Ae. aegypti* mosquitoes per visit varied monthly (Figure 2b), with a considerable jump after switching from standard CDC backpack aspirators to a handheld “Prokopack” aspirator [21]; the increased catch is indicated by the vertical dashed line in Figure 2. The monthly coefficient of variation exhibited overdispersion. Many months had a standard deviation greater than 1.5 the mean (Figure 2c). In space, there was similar heterogeneity in mosquito abundance (Figure 3). Across zones of the city, average adult female *Ae. aegypti* captured per visit varied from 0.24 to 0.71. Spatially, the coefficient of variation exhibited considerably larger variation than temporally, with values ranging from 1.2 to 17.1.

**Figure 2:**
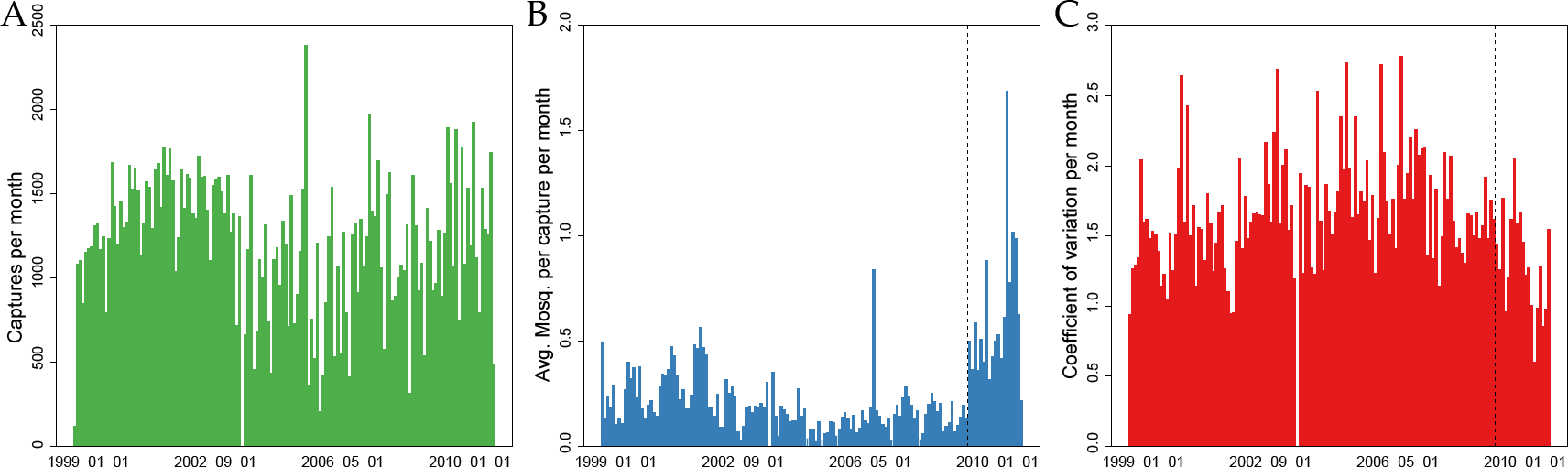
Collection effort by month, 1999-2010. Panel A: Number of collections per month are plotted. Panel B: Average number of female *Ae. aegypti* collected inside each house are plotted. Panel C: Coefficient of variation per month of number of female *Ae. aegypti* collected are plotted.

**Figure 3:**
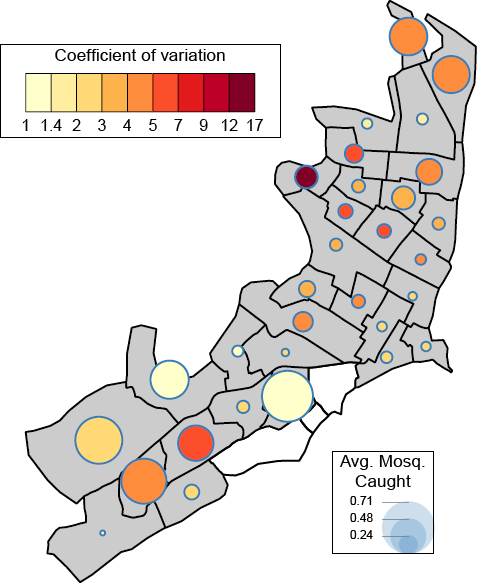
Coefficient of variation by space. By MoH Zone, the mean number of female *Ae. aegypti* caught per visit are plotted. Size of circle indicates average number caught and color of circle indicates coefficient of variation.

### Model development

#### Model structure and offset

From the base model (i.e., the model absent of any covariates other than the offset) overdispersion was clearly present (Poisson model AIC: 291417, negative binomial model AIC: 194781). The remainder of this section, therefore, focuses exclusively on the negative binomial version of the models. The over-dispersion parameter of the negative binomial base model, *θ*, was estimated to be 0.13. The fitted offset in an intercept and offset-only negative binomial model was 2.03 (1.99 – 2.08 95% CI). This value was used in all subsequent negative binomial models.

#### Full model overview

Due to the number of components of the mosquito abundance models, only key comparisons are detailed below. For a full description of all model comparisons, please see the supplementary information. The ‘full model’ structure is given in Figure 4. The fitted full model explained 10.3% of the deviance in the abundance data. In spite of the noise in the system that was unexplained, every component of the full model was found to be statistically significant (largest p-value was 2.5 x 10^−13^, 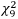 = 54.8, Table 1). The over-dispersion coefficient, *θ*, was estimated at its highest value of any model investigated as 0.152.

**Figure 4:**
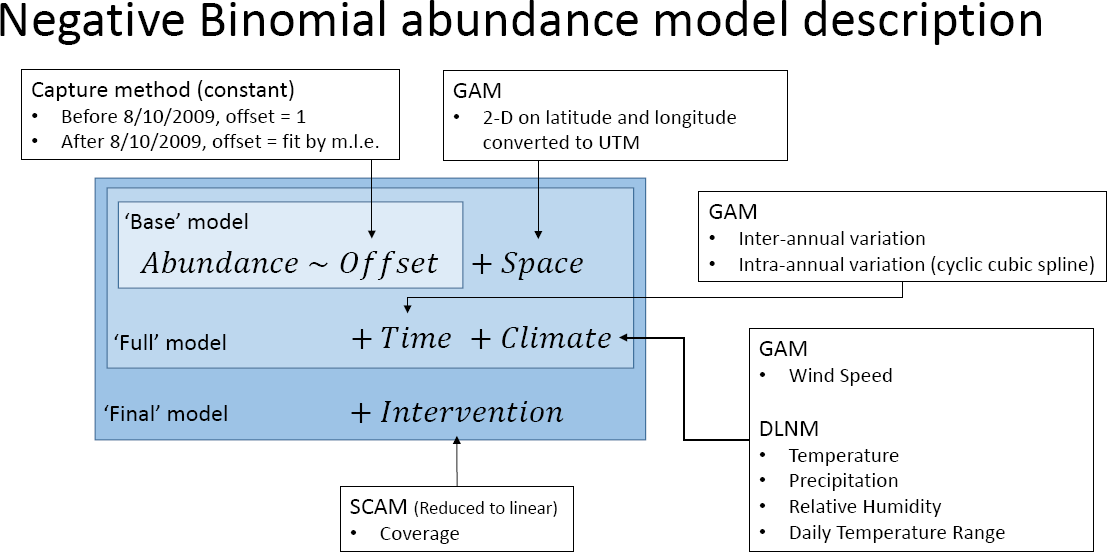
Model structure. Base model included only the offset due to collection method changes. The full model contained all spatial and climatological covariates. The final model then added the intervention data.

**Table 1:** Univariate and drop-one model results. Across each covariate, univariate and drop-one model statistics are listed.

|  | Univariate model versus base model |  |  |  |  | Drop-one model versus full model |  |  |  |  |
| --- | --- | --- | --- | --- | --- | --- | --- | --- | --- | --- |
|  | mle method |  |  | REML method |  | mle method |  |  | REML method |  |
| | $\Delta df$ | $\chi^2_{\Delta df}$ of LRT test | p-value | $\Delta AIC$ | $\theta$ | $\Delta df$ | $\chi^2_{\Delta df}$ of LRT test | p-value | $\Delta AIC$ | $\theta$ |
| **Space** | 28.1 | 1286 | <2e-16 | 1229 | 0.137 | 27.9 | 1154 | <2e-16 | 1098 | 0.147 |
| **Inter-annual** | 6.8 | 1756 | <2e-16 | 1738 | 0.139 | 7.5 | 976 | <2e-16 | 959 | 0.147 |
| **Intra-annual** | 8.9 | 730 | <2e-16 | 716 | 0.134 | 8.7 | 269 | <2e-16 | 256 | 0.151 |
| **Mean Temperature** | 9 | 1324 | <2e-16 | 1306 | 0.131 |  | - | - | - | - |
| **Maximum Temperature** | 9 | 1493 | <2e-16 | 1475 | 0.132 |  | - | - | - | - |
| **Minimum Temperature** | 9 | 1800 | <2e-16 | 1800 | 0.133 | 9 | 232 | <2e-16 | 214 | 0.151 |
| **Precipitation** | 9 | 1424 | <2e-16 | 1406 | 0.131 | 9 | 54.8 | 2.5e-13 | 37.3 | 0.152 |
| **Relative Humidity** | 9 | 1492 | <2e-16 | 1475 | 0.132 | 9 | 215 | <2e-16 | 198 | 0.151 |
| **Daily Temperature Range** | 9 | 1918 | <2e-16 | 1900 | 0.134 | 9 | 113 | <2e-16 | 94.3 | 0.152 |
| **Wind** | 2.9 | 3137 | <2e-16 | 3075 | 0.137 | 2.9 | 1726 | <2e-16 | 1721 | 0.152 |

For each component of the full model, two comparisons are detailed below. First, a univariate model evaluation comparing a model with no covariates (other than the offset) to a model with only the corresponding component. Second, a ‘drop-one’ model evaluation comparing a model with all components *except* the corresponding component to the full model with all components; results are summarized in Table 1. Finally, within each component subsection, the corresponding component’s contribution to the full model is discussed. Due to the use of generalized additive models (GAMs) within each biotic and abiotic covariate, it was difficult to provide the actual functional form of the relationships. The exact functional forms are available upon request from the authors.

#### Space

The addition of a general spatial covariate to the base model improved fit (ΔAIC = 1286). Space, however, was the second least significant addition to the base model. Conversely, dropping space from the full model caused the 2^nd^ largest reduction of fit (ΔAIC=1154), indicating that the variance explained by the spatial component was not explained by any other component. Again, all other covariates were aspatial. In the full model, the spatial component indicates a higher average abundance in the northeast and the southwest than other regions of the city (Figure 5a), with corresponding higher uncertainties in each of those locations (Figure 5b).

**Figure 5:**
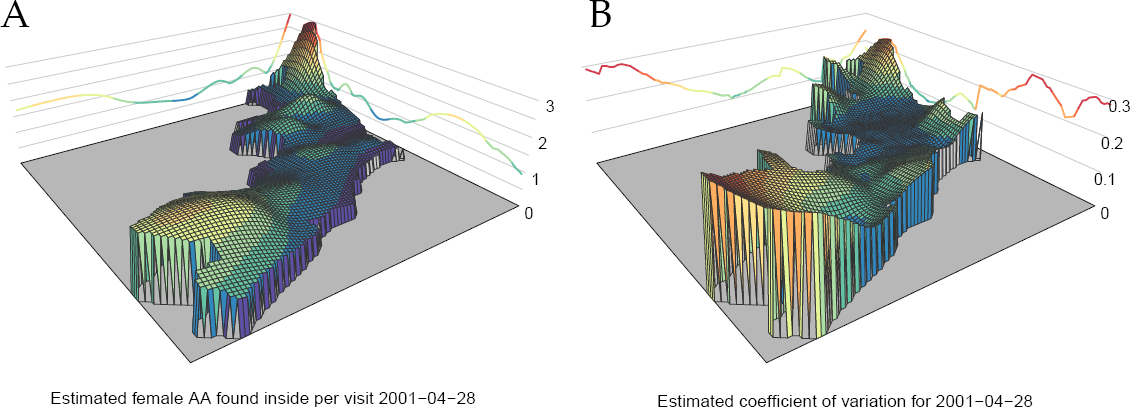
Fitted spatial variation in female *Ae. aegypti* abundance. Panel A: Estimated number of female *Ae. aegypti* that would be caught during a house-hold aspiration held on April 28^th^, 2001. The choice of this date was arbitrary but necessary to incorporate the climatological covariates. Panel B: Estimated coefficient of variation on number of mosquitoes for April 28^th^, 2001.

#### Inter-annual time

The univariate and ‘drop-one’ analysis for the inter-annual temporal term were conducted on the smooth versions of this component, but results are almost identical to one that was constant within years. Incorporating year-to-year variation resulted in the 4^th^ largest improvement on the base model (ΔAIC = 1756). In spite of the presence of all the temporally varying climate covariates, the inter-annual time component also resulted in the 3^rd^ largest reduction in model fit following its removal (ΔAIC=976). Within the full model, it appears that the early 2000s had considerably higher average mosquito abundance than later years, with 2006 being the lowest estimated population (Figure 6a).

**Figure 6:**
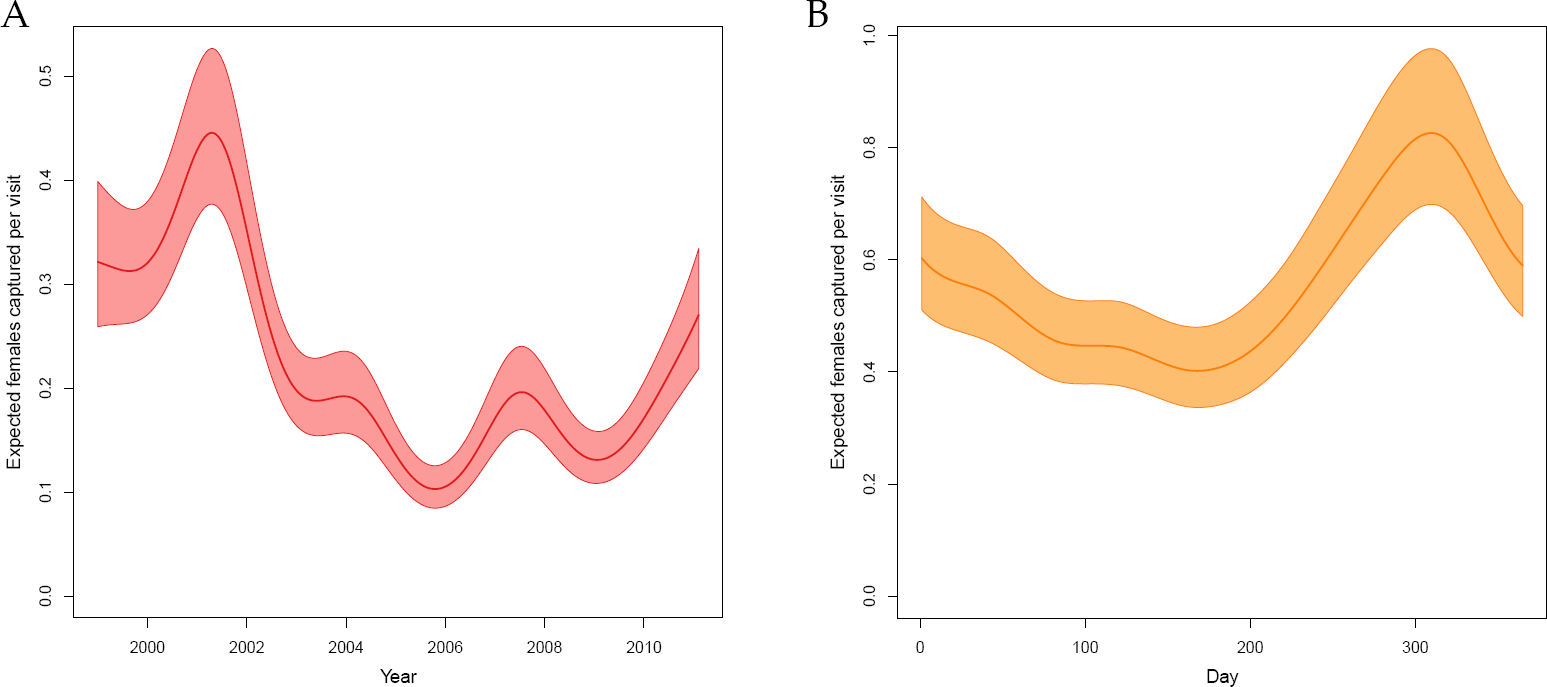
Fitted temporal variation in female *Ae. aegypti* abundance. Panel A: Estimated between-year (intra-annual) variation in the number of female Ac. *aegypti* that would be caught during a house-hold aspiration conducted at a home in the center of the city across the study period with 95% uncertainty. The choice of a home in the center of the city was arbitrary but necessary to incorporate the spatial variation. Panel B: Estimated within-year (inter-annual) variation in number of female *Ae. aegypti* that would be caught during a house-hold aspiration conducted at a home in the center of the city in 2001 with 95% uncertainty. This choice was arbitrary but necessary to incorporate intra-annual variation.

#### Intra-annual time

Seasonality alone performed relatively poorly, improving the model by the least amount of any component (ΔAIC=730). The seasonal term still appeared to explain variation above and beyond the climate terms as indicated by a significant, but relatively moderate, reduction in fit of the full model when the seasonal term was removed (ΔAIC=269). In the full model, there was an average low mosquito abundance in June and July and a peak around October (Figure 6b).

#### Weather

Each of the climate components significantly improved fit compared to the base model (Table 1). Among the temperature covariates, the model that included minimum daily temperature performed better than when mean or maximum daily temperature were added to the base model (ΔAIC=1800 versus ΔAIC=1324 and ΔAIC=1493, respectively). As such, it was retained for further evaluation. Across all components, wind on the day of collection had the largest overall impact on the base model (ΔAIC=3137).

The importance of wind was again found in the ‘drop-one’ analysis with the full model deteriorating the most following removal of wind (ΔAIC=1721). The least important component was precipitation, although it was still strongly significant (p-value of 1.3 x 10^−13^). Daily temperature range, which was 2^nd^ most important in the univariate analysis (ΔAIC=1918), was 2^nd^ least detrimental to drop from the full model (ΔAIC=113).

The impact of each climate variable in concert with all other components and in isolation is discussed in the supplementary information and plotted in Figures S1-S5 and Figures S6-S9, respectively.

### Assessing vector control

Inclusion of the intervention term further improved fit compared to the full model (ΔAIC=144). Inspection of the fitted shape-constrained additive model (SCAM) revealed that the smooth for control was linear (edf = 1). To simplify the model, we reverted to the GAM version of the model with the intervention term entering linearly.

The negative binomial model has a log link, so terms that enter ‘linearly’ in the model relate to the response exponentially. The coefficient for the intervention term was found to be −1.1 (standard error 0.09). Figure 7 shows the predicted impact of various levels of intervention on a hypothetical home where, in the absence of control (or “dose response curve”), 0.45 female *Ae. aegypti* mosquitoes were expected to be collected. If all the homes were sprayed across the zone (i.e., the intervention coverage was 1), we would expect to see a 67% reduction to 0.15 mosquitoes. If half of the homes in a zone were sprayed, and thus the ‘intervention coverage’ was 0.5, we would expect to see a reduction to 0.26 mosquitoes in that same hypothetical home; a 43% reduction.

**Figure 7:**
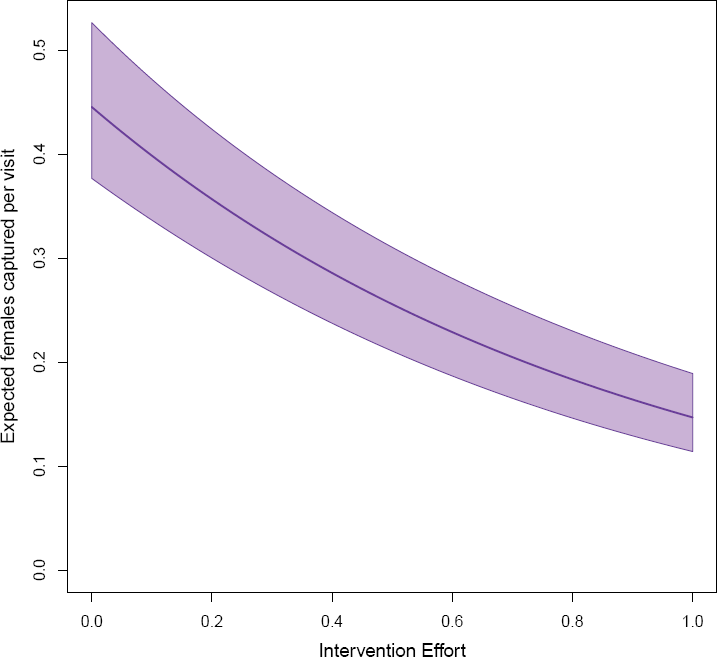
Dose-response curve for the Ministry of Health space-spray intervention. Expected changes in number of female *Ae. aegypti* captured with 95% uncertainty as a function of impact of various levels of intervention effort on a home in the center of the city measured on April 28^th^, 2001. These choices (home in the center of the city, the date of April 28^th^, 2001) are all arbitrary but necessary to account for spatial and temporal variation.

## Discussion

Our aims were two-fold. First, to identify the dose-response curve between neighborhood-level spraying interventions and neighborhood *Ae. aegypti* population abundances. To achieve in the face of substantial spatio-temporal variation in mosquito population abundances and imperfectly sampled mosquito populations, we had to achieve our second goal, to develop a mosquito abundance-modeling framework. Our model had to account for both biotic and abiotic drivers of mosquito population dynamics and provide counter-factual estimates of what mosquito populations might have looked like had spraying not been conducted. By applying our model to Iquitos, Peru, we identified interpretable relationships between climatic covariates and mosquito populations, as well as, site-specific spatial variation, with some regions of the city appearing to over- or under-produce mosquitoes consistently through the study period. When we then added an analysis of the existing space-spraying campaigns conducted by the Iquitos Ministry of Health, we found a diminishing return dose-response curve where, under optimal settings, we would expect to see a 67% reduction in female *Ae. aegypti* abundance from pre-intervention levels. This means that even a perfectly enacted Ministry of Health space-spraying campaign would not be adequate to reduce mosquito populations to zero.

Univariate analyses conducted on each climatological covariate revealed and confirmed relationships between key features of local climate and mosquito population dynamics. For temperature, precipitation, relative humidity, and daily temperature range, the effect of climate on mosquito abundance was not isolated to a particular lagged covariate. As expected, weather that directly influenced mosquito dynamics a few generations in the past indirectly influenced the number of mosquitoes collected on any given day. These results, and results from our full model analyses, highlight the importance of measuring daily weather when estimating mosquito abundance baselines. Our modeling approach continuously integrates data from the day of capture back through 30 days in the past, and we found significantly different influences of these weather variables at various lags. Factors that are rarely measured, such as maximum wind speed, were found to be strongly significant and should be considered for future entomological studies on *Ae. aegypti* abundance. Previous analysis also found that winds may influence the spatial patterns of dengue transmission in Iquitos [22]. It was not possible for us to assess the reason why high wind speeds was associated with higher capture rates in homes, but a positive relationship between wind speed and aspirator collection of *Ae. aegypti* is a plausible explanation. Windy conditions may increase the probability that mosquito sequester in indoor locations that are less effectively sampled than during more calm days.

Our analysis of neighborhood spraying accommodated a variety of potential functional forms for the dose-response curve, yet the model-selection process yielded the simple, exponential decay as the approach with more support in explaining the data. Perhaps it is unsurprising that we found a diminishing return relationship between spray effort and mosquito reduction, and it may be that the true optimal effectiveness of the intervention we analyzed was a 67% reduction in adult mosquitoes. Importantly, this analysis ignored the likely super-linear increase in costs that would be associated with increasing effort. In the absence of an analysis that incorporates cost explicitly, it is difficult to recommend to a ministry of health that complete coverage is the only goal. Even a 50% coverage still achieved a reduction of mosquito abundance by 43%, and this would likely be considerably less costly to enact.

Although our study is based on a large data set, analysis of over 175,000 individual home visits, we were not able to overcome all of the inherent limitations associated with mosquito abundance estimation. *Aedes aegypti* have cryptic landing and larval development sites [23, 24]. Aspiration, even when conducted by a team of trained professionals, is an imperfect method to fully survey a house. Our *Ae. aegypti* abundance estimates for Iquitos underrepresent the true number of mosquitoes within the city. We similarly lacked spatially varying climate data. Climate at the local airport may not represent conditions in the houses where mosquitoes were collected. Microclimates that vary at the same scales as neighborhood blocks have been shown to influence *Ae. aegypti* ecology [25] and biting habits [26]. The metric we used to assess the impact of intervention for our analysis is an imperfect match to account for exactly which houses did and did not receive an intervention on which days. Although the data available in Iquitos was very detailed, even in this setting, more careful collection of what was done when and where would improve the ability to disentangle the effects of interventions from other factors. As efforts to control arbovirus transmission continue, cataloging detailed intervention activities should be a priority so that more of the work being done can be used to inform future control approaches.

A significant limitation of our analyses was that it does not attempt to link mosquito abundance with virus transmission intensity. Mosquito abundance is only one of many spatio-temporally varying drivers of mosquito-borne pathogen transmission. Dengue virus transmission intensity in Iquitos varies from year to year [27]. An analysis of human dengue cases in Iquitos indicated that the incidence of people with symptomatic disease was negatively correlated to insecticide space spraying efforts only when the intervention was conducted early in the virus transmission season [28]. Our analyses indicate that interventions should have had an impact on mosquito populations independent of season. Spatially, there is substantial evidence that human movement is a strong driver of the spatial patterns of DENV transmission [29, 30]. It is unlikely, therefore, that targeting an intervention based solely on a model of mosquito abundance would capture all of the places contributing most to transmission. Although we expect that reducing mosquito populations will correlate with a reduction in transmission risk in many instances, our modeling framework is intended to serve as a part of a greater integrated modeling approach that attempts to optimally target interventions given adequate knowledge of the ecology of the mosquito vector, its human hosts, and the pathogen.

Results from our analyses demonstrate that, even in the face of considerable unexplained variation, drivers of mosquito abundance can be identified and even a weak correlate for intervention effect size can identify a significant control impact. We provide a statistical framework that can be used to establish a mosquito abundance baseline in space and time. With a baseline, we are able to assess the impact of mosquito control on abundance by estimating the difference between the observed number of mosquitoes caught and the expected number caught if there was no intervention. Understanding the spatial scales of mosquito heterogeneity and mosquito control is essential for accurately accounting for variation in transmission risk during the design and execution of vaccine and vector control trials [31]. Likewise, an improved understanding of heterogeneity can inform the application of vector population reduction (e.g., genetically engineered mosquitoes [32]) and population replacement (e.g., *Wolbachia*) programs [33]. In the absence of reliable and accurate estimates of mosquito abundance, vector control programs will likely miscalculate, overestimate or underestimate, the required initial intensity and effect size of their intervention. The approaches described in our framework are intended to help address this gap by identifying site-specific drivers of mosquito abundance across different spatial and temporal scales in a diversity of ecological and epidemiological contexts.

## Materials and Methods

### Data

Ethics Statements Entomological survey data were collected under protocols approved by Institutional Review Boards at the University of California at Davis, Tulane University, Emory University, San Diego State University, Liverpool School of Tropical Medicine, London School of Tropical Medicine and Hygiene, Peruvian National Institute of Health, Naval Medical Research Center, Bethesda, and U.S. Naval Medical Research Unit No. 6. The latter included Peruvian representation, in compliance with all US Federal and Peruvian regulations governing the protection of human subjects. All protocols were reviewed and approved by the Loreto Regional Health Department, which oversees health research in Iquitos.

Mosquito Collection Data: Adult *Ae. aegypti* were collected in Iquitos households as part of ongoing prospective cohort studies (Table S1). Sampling schemes varied with different study designs and were concentrated in the northern two districts of Iquitos (Maynas and Punchana), with the exception of some non-residential sites in 2004-2005, one October 2006 survey, and a 2009-2011 randomized cluster study carried out in the southern district of San Juan. Daily entomological surveys (methodology described in detail by Morrison et al. and Schneider et al. [19, 34]) were carried out Monday through Friday, between the hours of 0700 and 1300, by the same 6 two-person collection teams from 28 January 1999 through 8 February 2011. Briefly, each survey consisted of a short questionnaire, inspection of the household for water-holding containers, and collection of adult mosquitoes initially using a backpack aspirator (Clark) and then, beginning in June 2009, a Prokopack Aspirator [21].

### Mosquito control methods

Ministry of Health Control (MOH) Activities were irregular prior to 2002, when the first well-organized city-wide emergency mosquito control campaigns were initiated. Starting in 2002, stable entomological (both surveillance and control) teams distributed the larvicide temephos to wet containers focusing on permanent or semi-permanent water or abandoned containers (i.e., tires and appliances) and when possible destruction of smaller containers. In theory, these teams visited neighborhoods at approximately 3-month intervals, but coverage rates were highly variable between the 34 zones designated by the MOH. The first well organized and comprehensive city-wide campaign was carried out between 23 October 2002 and 24 January 2003. Focal application of temephos was followed by 3 weekly cycles of door to door intradomicile space spray with Deltamethrin (0.25%). Another large disease outbreak led to a similar city-wide campaign initiated in December 2004. Smaller more spatially focused interventions, which were administered with backpack sprayers, occurred in February 2003, October 2003, January 2005, February 2006, and November 2006. During these interventions the chemicals used varied between Alphacypermethrin and Deltamethrin. Adulticiding was not carried out in 2007. More geographically limited activities were carried out in the first and the last trimester of 2008 and the first trimesters of 2009 and 2010 with either alphacypermetrin or cypermetrin. Spray schedules were developed by MOH zone. Coverage levels were not recorded.

Mosquito collections were carried out in the context of three vector intervention trials. The first was carried out on 12 city-blocks between February and December 2005 and included a onetime application of a residual formulation of lambdacyhaltrin followed by treatment of larval habitats with BTI at monthly intervals. In the second, from March 2007 to December 2007 a control trial based on clusters of 5 city blocks in each of 10 MOH zones was treated with a different larvicide; either temephos or pyriproxyfen. This study did not include adulticiding. In the third, from September 2009 through 2010 a randomized cluster trial evaluated Deltametrin treated curtains in the San Juan District. Treated curtains were distributed among houses in 10 clusters of 1-3 city blocks each. The MOH interventions were done independent of whether a home was within a control or a treatment area.

Climate Data: Daily climate data for Iquitos was acquired and processed as in Stoddard et al [28]. Briefly, daily measures of temperature (mean, minimum, and maximum), precipitation, maximum wind speed, and dew point were attainted from a US National Oceanic and Atmospheric Administration (NOAA) weather station located at the Iquitos International Airport. Weather data was used to generate daily temperature range and relative humidity. Because there was only one NOAA station in Iquitos, we had no measurement of spatial variation in climate covariates in our analyses. We applied the same weather data to all houses identically.

#### Model development

Poisson regression: From first principles, count data (such as the number of mosquitoes captured inn a given house on a given day) can be modeled using a Poisson distribution. Given the presence of covariates that could drive changes in the expected number of mosquitoes caught per visit across space and time, this extends to a Poisson regression model. A common difficulty with count data is that the variance is often considerably larger than the mean; i.e., over-dispersion. When over-dispersion is present, a Poisson model is no longer appropriate. There are multiple alternative options, including zero-inflated Poisson, negative-binomial, and zero-inflated negative-binomial. Throughout, when possible, all options were fit. To assess the relationship between the variance and the mean for the regression models, we plotted a version of the coefficient of variation that was more appropriate for skewed data [35]:

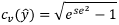

where *se* is the standard error of prediction 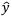 in log space; i.e., untransformed back to the natural scale.

The final model structure is given in Figure 4. Below, the model structure is broken down in its constituent parts with each part describing the covariates used, as well as the way in which they enter the full model.

Offset: On 1 August 2009, collections switched from backpack aspirators to Prokopack aspirators. Previous analysis indicated that the Prokopack aspirators doubled the number of mosquitoes collected and, as such, a different offset was used after this date. Before that date, the offset was set to 1, and the offset afterwards was fit by maximizing the likelihood of the base model. The confidence interval for the offset was calculated using the profile likelihood technique, which inverts the single parameter likelihood ratio test (Figure S12).

Space and Time: Simple summary statistics and graphical summaries (Figure 2, Figure 3) were consistent with results from other studies [13,14], which showed that there was spatial and temporal variation in mosquito abundance in Iquitos. To account for this variation while avoiding enforcing any particular functional form to these relationships, flexible splines were incorporated for both abiotic spatial and abiotic temporal drivers of variation. For space, a twodimensional smooth incorporated the exact location of each home, while two one-dimensional smooths incorporated inter- and intra-annual temporal variation. These smooths were then fit within the framework of a generalized additive model (GAM). We implemented these models using the mgcv package [36] in R.

Climate: The influence of each climatological covariate on *Ae. aegypti* population dynamics and the probability of collecting adult *Ae. aegypti* is complex. Because both the underlying dynamics and the effects of the observation process are estimated simultaneously within the count regressions, particular functional forms of these relationships do not necessarily follow directly from those observed in controlled experiments. For example, the relationship between temperature and mosquito survival is generally clear. At both low and high temperatures, mosquitoes are less likely to survive than at moderate temperatures. In field settings, however, temperature fluctuates throughout each day and from day to day. Temperature on any given day accounts for only a part of the total contribution of “temperature” on the eventual number of mosquitoes caught. Moreover, because field conditions are not a controlled experiment, unmeasured covariates that occur collinearly, such as collector bias or temperature may alter the fitted relationship between temperature and mosquito abundance. Typically, lagged covariates account for the expected effect of past weather on current observations, with the length of the lag either estimated or fixed based on previous studies. This approach ignores the cumulative effect of all past weather on eventual observations, with at most only a few lagged covariates attempting to capture the entirety of the past.

A recently developed modeling tool, distributed lag models [37], estimates the relative weights that each moment in the past exerts on the present, integrating across the past. Because the relationships between climate and mosquito population dynamics are frequently non-linear, we incorporated non-linear functional forms (i.e., splines) to account for both the relationship between the covariate and the response as well as the lag weights. The resulting models for the climate covariates were distributed lag non-linear models (DLNMs). The exception in this analysis was that maximum wind speed was not assumed to influence population dynamics, only the probability of capturing mosquitoes on the given day. As such, maximum wind speed was incorporated with no lag and was included in the models as a one-dimensional smooth. Due to the number of parameters and covariates in this exercise, the type of smooth as well as the number of knots in lag space and covariate space for each DLNM was selected to be the same. Specifically, natural splines were used throughout for climate covariates, and lag and covariate space degrees of freedom were set to 3. This allowed for a moderate amount of flexibility within the lag and covariate relationships. All maximum lags were set at 30 days to avoid overfitting. We implemented the DLNM component of our models within the dlnm package [37] in R.

Intervention: One of the drawbacks from the flexibility of GAMs is that they may select for biologically impossible relationships. Here, ‘intervention effort’ represented the number of homes within a zone that were sprayed. While a high value of this covariate for a given home did not imply that this home actually received treatment, it did imply that more of the area around that home was treated. Thus, for this analysis, we enforced a monotonically nonincreasing relationship between ‘intervention effort’ and abundance. The shape of the relationship was allowed to be non-linear, but as effort increased, we forced abundance to not increase. If the data only supported a positive relationship between effort and abundance, the result of our assumption would be a fitted relationship that was a flat line. Each time we incorporated the shape-constrained additive model, we did so within the scam package [38] in R.

Model fitting and selection: Before model fitting, each variable was centered and standardized. The final models presented were fit, when possible, using restricted maximum likelihood (REML), because this approach reduces bias and over-fitting of the smooth splines [39]. Models were also fit maximizing the likelihood as well as generalized cross-validation (GCV). Model selection within the GAM framework can be approached in several ways. Within these analyses, since we focused mostly on REML models, we predominantly used Akaike information criterion (AIC), but every comparison selection that used GCV or maximizing likelihoods would have made the same decision. For all results in the main text, we give results both in terms of ΔAIC for REML models and likelihood ratio tests for maximum likelihood models. The full model was built in the following order; after each step, if necessary, the model was reduced. First, spatial covariates were added. Second, the non-climate temporal covariates were added. Third, the climate covariates were added. The “full” model was then reduced by sequentially dropping each covariate and assessing AIC scores. After reduction, the control covariate was added to the model. The final model was again reduced to assess the shape of the control effect. All calculations were done with R 3.3.1 [40].

#### Assessing vector control

Mosquito control data documented which zone of the city was covered, the number of homes within the zone, and the number of homes that received treatment. There was no information on exactly which homes within a zone were sprayed. Most control efforts consisted of three sequential applications of insecticide in the same zone over the course of one to two weeks. Across sequential applications, there was no indication of whether the same subset of homes received all, some, or none of the treatments. Consequently, in our analyses we were not able to incorporate ‘control’ at the household level. Instead, we applied vector control to entire zones of the city simultaneously. For each home in a treated zone, we defined the ‘intervention effort’ as the percent of the homes that were treated. Although homes that were not treated have imputed non-zero ‘intervention effort’ scores, this covariate had the desired properties such that if the score was equal to 0, we were certain that all homes in a zone were not treated, and if it was equal to 1, we were certain that all homes in the zone were treated.

Results from recent neighborhood studies in Iquitos indicate that *Ae. aegypti* populations return to baseline within 3 weeks of an indoor insecticide space spraying application (ACM unpublished data). As a result, we not only imputed ‘intervention efforts’ to each home in a zone during the period of spraying, but we also extended the spray “effect” to 3 weeks after the end date of spraying. When a zone received multiple sequential interventions, the ‘intervention efforts’ did multiply. Rather, if two interventions in the same zone occurred within 3 weeks of each other, the ‘intervention effort’ for each round of spraying was calculated separately for each day and the maximum effort across rounds was retained (Figure S10).

## Supporting Information

Results from the base model confirmed that there was statistically significant over-dispersion (Poisson model AIC: 291417, negative binomial model AIC: 194781). The base model (with only the offset as a covariate) performed poorly in some cases, with the Poisson model explaining 0% of the deviance and the negative binomial model explaining 3.9%. Including an additional twodimensional smooth representing abiotic/unexplained spatial variation significantly improved fit (Poisson model AIC: 287633, negative binomial model AIC: 196340). The addition of the two abiotic temporal smooths further increased fit significantly (Poisson model AIC: 280361, negative binomial model AIC: 193940). For comparison purposes, a model without the spatial smooth, but with the two temporal smooths was fit as well. The model with only the temporal variation accounted for performed better that the model with the spatial smooth (Poisson model AIC: 284082 vs 287633, negative binomial model AIC: 195178 vs 196340).

Among the temperature covariates, the model that included minimum temperature performed better than when mean or maximum temperature were included (negative binomial AIC of 192539 versus 192588 and 192857, respectively). The full model increased the deviance explained considerably (5.6% and 10.3% respectively). The ‘drop one’ test did not identify any covariates that should be dropped (Table 1). Judged by the change in AIC, the least influential covariate for the negative binomial model was daily temperature range, with a drop in fit corresponding to an increase in AIC of 95. The most influential covariate was maximum wind speed, with a drop in fit corresponding to an increase in AIC of 1721.

The inclusion of the control covariate further improved the fit. Inspection of the fitted SCAM model revealed that the smooth for control effort was found to be linear. To simplify the model, we reverted to the GAM model with control effort entering linearly. For the negative binomial model, the final model with control effort linearly added to the reduced model, improving AIC by a further 144 points.

To show the relationship between any given covariate and the response for all plots from the final model, we held all other covariates constant. Unless otherwise indicated, all plots represent the abundance in the center of the city on 28 April 2001, with all climatic drivers set to their mean and no control assumed. This date was chosen because it was estimated to have the highest expected abundance.

The final model clearly captured variation in space (Figure 5, Figure S11). On 28 April 2001, the range of expected mosquitoes found per visit varied from almost 0.5 up to 3 with the highest expected abundance in the far north. There was a greater coefficient of variation on the periphery of the city, with a noticeable increase in the north where the greatest expected mosquito counts occurred. Within-year variation (Fig 6b) showed clear seasonality, with the most mosquitoes expected late in the year and the fewest in June and July. These dates matched the timing of the dry and wet seasons in Iquitos. After controlling for the influence of climate, 2002 was the year with the lowest mosquito abundance (Figure 6a). Wind speed had a strongly non-linear relationship with abundance, with higher counts typically observed on windier days (Figure S1).

The DLNM relationships were each plotted in a similar manner to each other. In the left panel, the z-axis shows the multiplicative effect of various combinations of lag and climate value. The lag axis shows the effect of climate up to 30 days in the past, while the front axis shows the range of values of the covariate. For each of these figures, interpretation requires envisioning what the value of the covariate was over the course of the past 30 days and then averaging over this curve on the surface, somewhat analogous to line integrals. The predicted values were all centered on the mean climate value, so each surface is constant at 1 for the mean climate value across lags. For the remaining 6 panels, slices of the surface were taken and plotted with 95% confidence intervals. The center panels display the relationship across different lags between the climate variable at the lowest, middle, and highest values observed and expected mosquito counts. These represent the expected relationship between the climatic driver and mosquito abundance when the climatic driver was held constant at that value over the past 30 days. The right panels slice the DLNM surface across lags of 0, 10, and 20 days. These represent the relationship between the climatic driver and mosquito abundance at the moment of capture as well as 10 and 20 days in the past. Again, for the right panel, the relationship was plotted using the mean climatic value as the reference value. These relationships go through 1 with a vanishing confidence interval at that point.

For the full model, it is important to note that identified relationships are different than in univariate climate models due to collinearity in the climate covariates. The variation captured by each climatological driver must be interpreted in the presence of all other drivers. For minimum temperature (Figure S2), there was an almost quadratic relationship between lag and response for low temperatures and essentially no multiplicative effect for high temperatures across lags. Precipitation was skewed right with the mean value of 0.96. As such, the reference value was close to the lowest observed value and thus the top center panel of Figure S3 shows no multiplicative effect by construction. Conversely, large rainfall a month in the past resulted in an increased expected count. Similarly, large rainfalls 2 weeks before capture reduced expected abundance. For daily temperature range (Figure S4), large variation in daily temperature 30 days in the past increased the expected number of mosquitoes caught. For low variations in daily temperature, there was only a small effect. Recall that in the full model, daily temperature range was the covariate closest to being dropped. The relationship between relative humidity (Figure S5), time, and expected mosquitoes caught was complex. Iquitos is in the Amazon Basin where the environment is quite humid throughout the year; relative humidity ranges from 61% to almost 100%. ‘Moderate’ humidity of 84.6% was used as the baseline because this was the mean estimated daily relative humidity and thus had no multiplicative effect on mosquito abundance. Either ‘drier’ or wetter environments may increase or decrease abundance depending on when in the development stage of the mosquito the weather changes.

For the univariate climate models, relationships changed between each driver and response. These models were fit with all non-climatic drivers and then only a single climatic driver at a time. This allowed for a more direct interpretation of the climate variable’s influence on mosquito abundance. When minimum temperature and abundance were assessed in isolation (Figure S6), the relationship more closely matched the expectation derived from survival studies [41]. At both high and low temperatures, abundance was expected to decrease relative to moderate temperatures. That relationship was mostly constant in lag space, with only substantial curvature at extremely low temperatures, where expected counts were already decreased in all cases. The relationship between precipitation and abundance also changed from the full model. At a lag of 0, there was a mostly monotonically increasing relationship between precipitation and expected abundance, while at earlier lags this relationship was reduced (Figure S7). Daily temperature’s relationship to abundance switched in some cases (Figure S8) relative to the multivariate analysis with high variation in daily temperature a month in the past, resulting in a large decrease in expected mosquito abundance. For relative humidity (Figure S9), at moderate lags the relationship followed expectation, with higher humidity resulting in higher abundance. At no lag, the relationship was quadratic, with a decrease in expected abundance at both high and low relative humidity levels. In this context, ‘low’ is still 61% relative humidity.

## Figures

**Figure S1:**
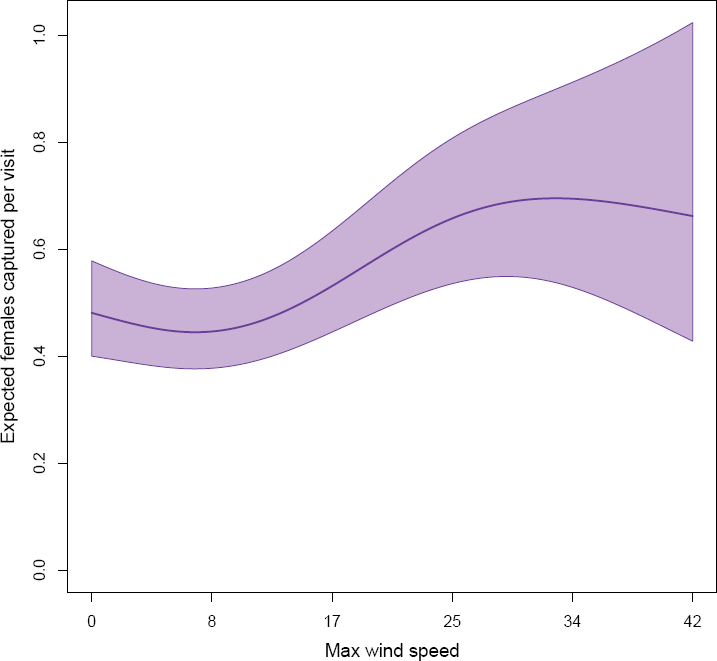
Effect of max wind speed on mosquito abundance. Expected number of female *Ae. aegypti* captured with 95% uncertainty as a function of wind speed on a home in the center of the city measured on April 28^th^, 2001. These choices (home in the center of the city, the date of April 28^th^, 2001) are all arbitrary but necessary to produce estimated counts that account for spatial and temporal variation.

**Figure S2:**
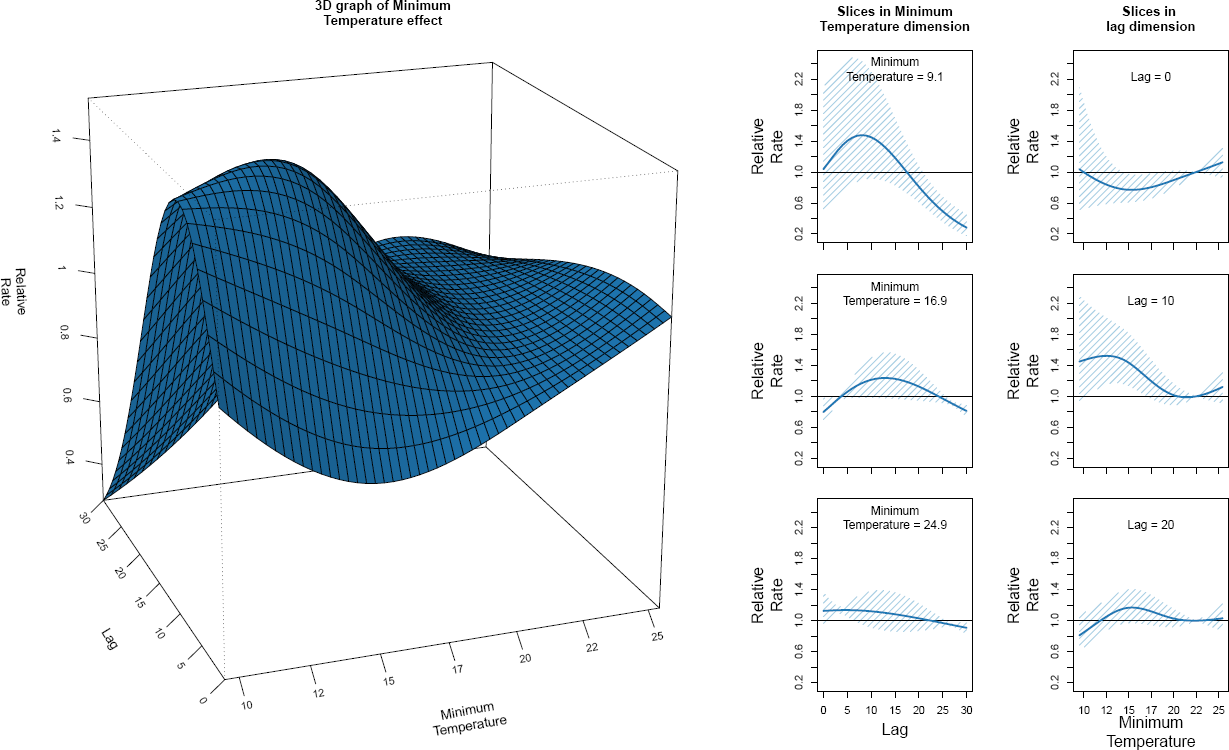
Effect of lagged daily minimum temperature on mosquito abundance in the full model. Panel A: The relationship between minimum temperature (x-axis), lag up to 30 days (y-axis), and relative rate of number of mosquitoes caught (z-axis) is plotted. Panel B, D, and F plot slices of the surface along the temperature axis with corresponding uncertainty. Panel C, E, and G plot slices of the surface along the lag axis with corresponding uncertainty.

**Figure S3:**
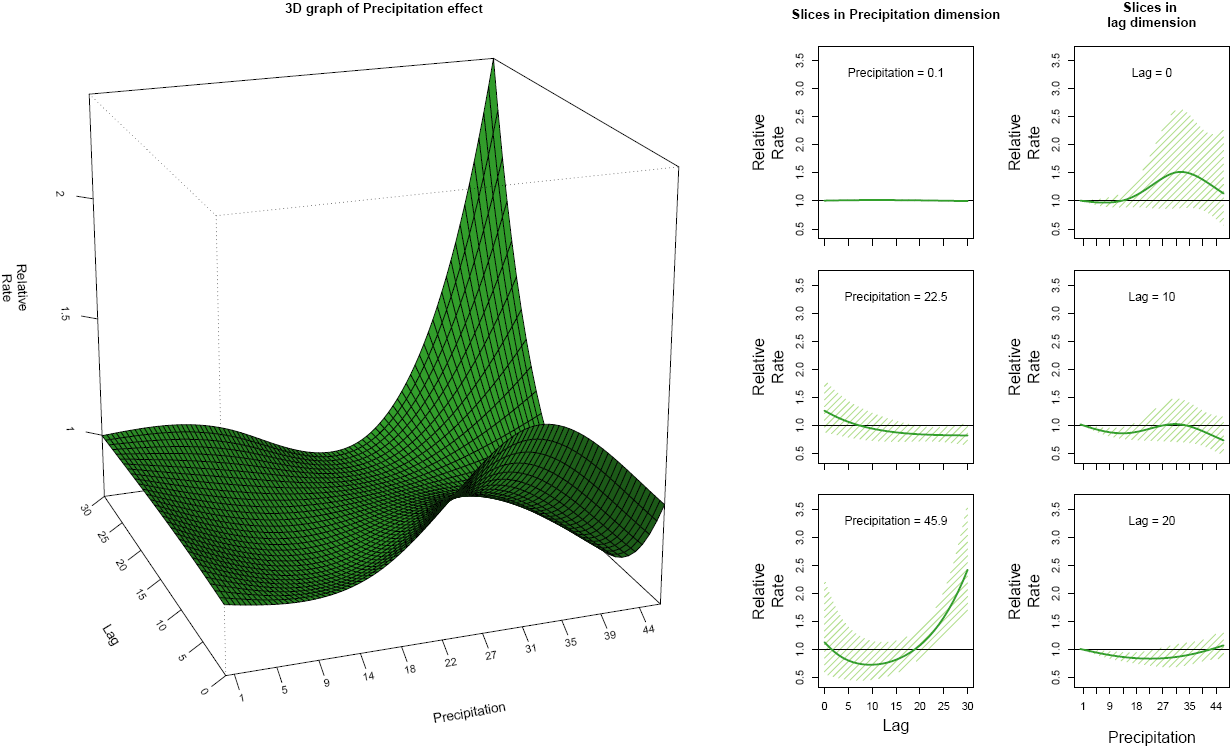
Effect of lagged daily precipitation on mosquito abundance in the full model. Panel A: The relationship between precipitation (x-axis), lag up to 30 days (y-axis), and relative rate of number of mosquitoes caught (z-axis) is plotted. Panel B, D, and F plot slices of the surface along the temperature axis with corresponding uncertainty. Panel C, E, and G plot slices of the surface along the lag axis with corresponding uncertainty.

**Figure S4:**
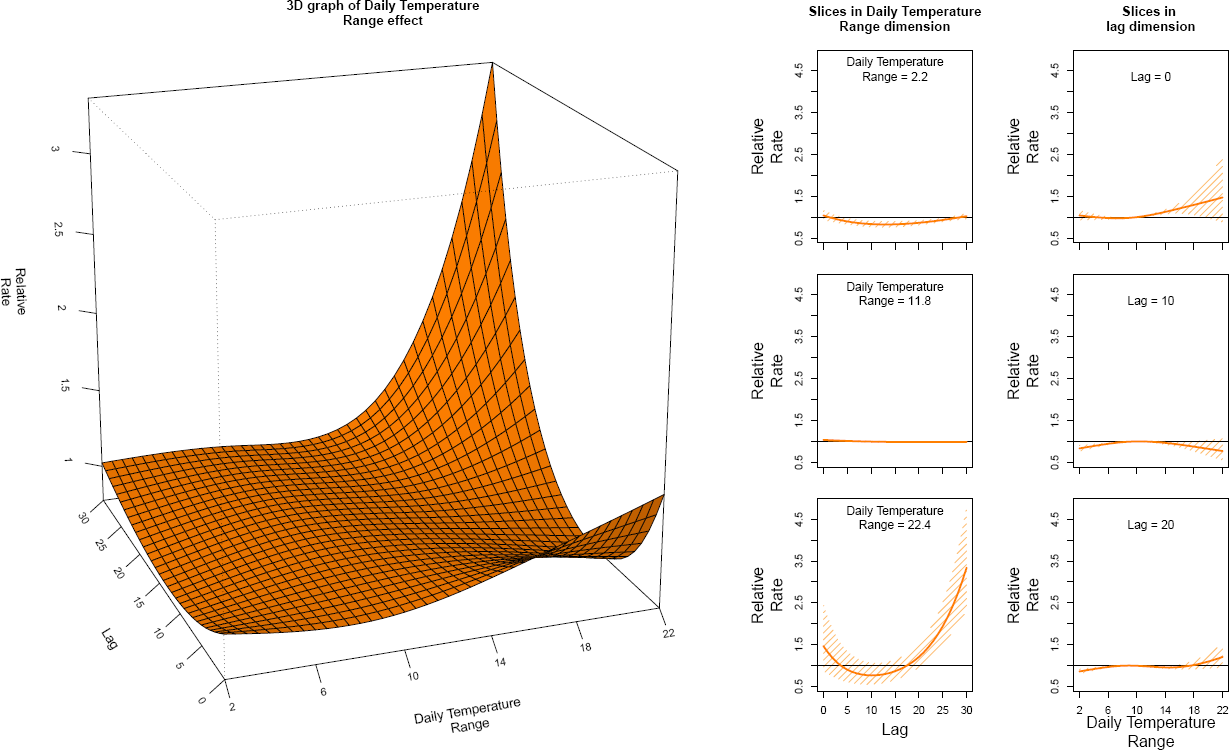
Effect of lagged daily temperature range on mosquito abundance in the full model. Panel A: The relationship between daily temperature range (x-axis), lag up to 30 days (y-axis), and relative rate of number of mosquitoes caught (z-axis) is plotted. Panel B, D, and F plot slices of the surface along the temperature axis with corresponding uncertainty. Panel C, E, and G plot slices of the surface along the lag axis with corresponding uncertainty.

**Figure S5:**
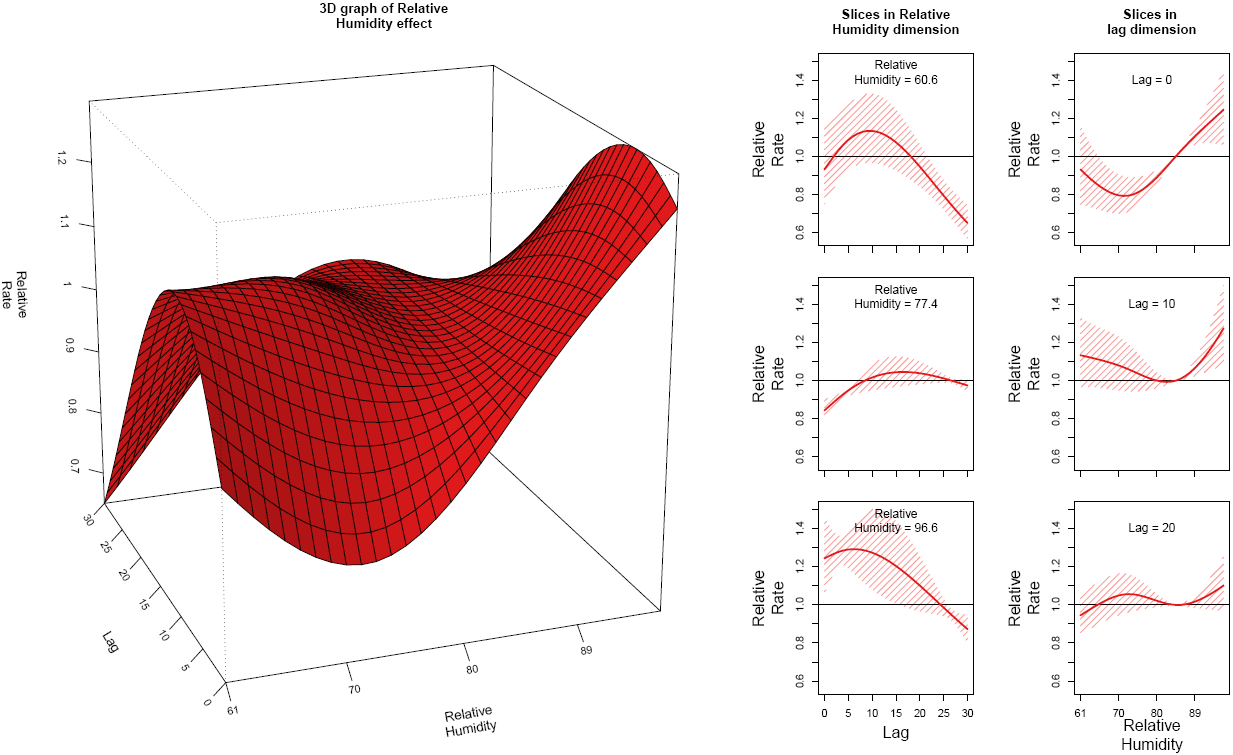
Effect of lagged daily relative humidity on mosquito abundance in the full model. Panel A: The relationship between relative humidity (x-axis), lag up to 30 days (y-axis), and relative rate of number of mosquitoes caught (z-axis) is plotted. Panel B, D, and F plot slices of the surface along the temperature axis with corresponding uncertainty. Panel C, E, and G plot slices of the surface along the lag axis with corresponding uncertainty.

**Figure S6:**
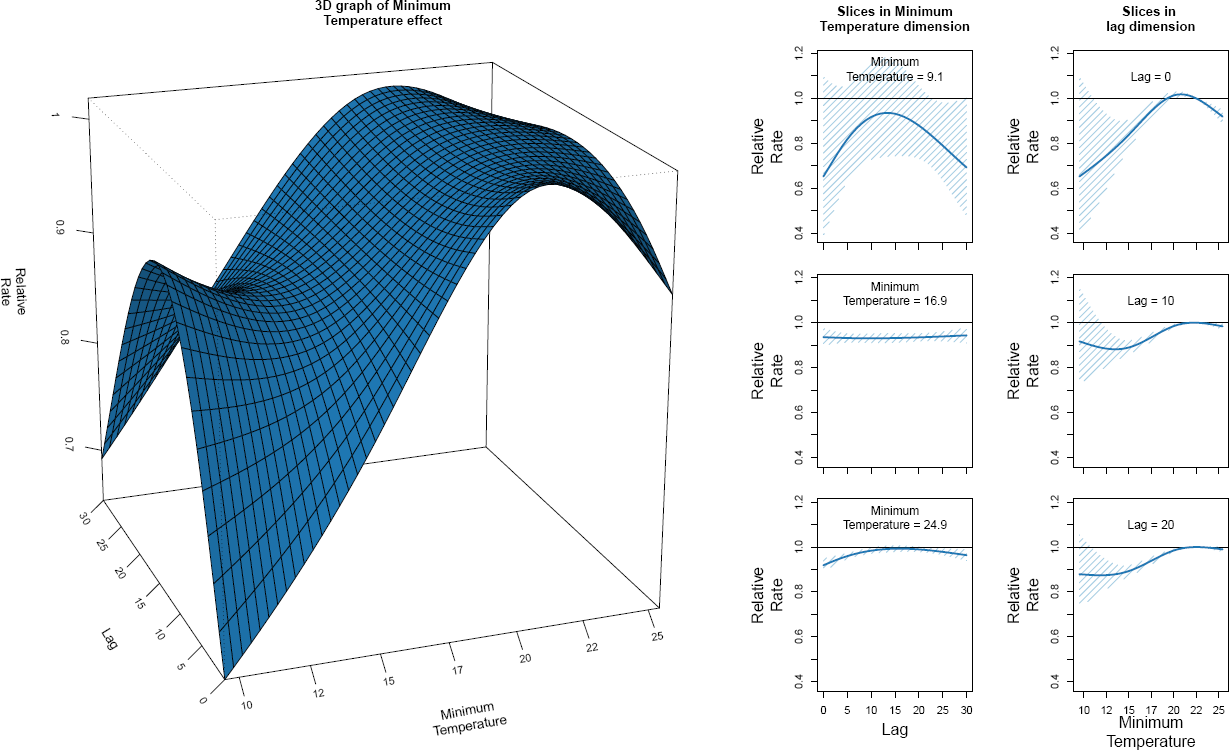
Effect of lagged daily minimum temperature on mosquito abundance in a univariate model. Panel A: The relationship between minimum temperature (x-axis), lag up to 30 days (y-axis), and relative rate of number of mosquitoes caught (z-axis) is plotted. Panel B, D, and F plot slices of the surface along the temperature axis with corresponding uncertainty. Panel C, E, and G plot slices of the surface along the lag axis with corresponding uncertainty.

**Figure S7:**
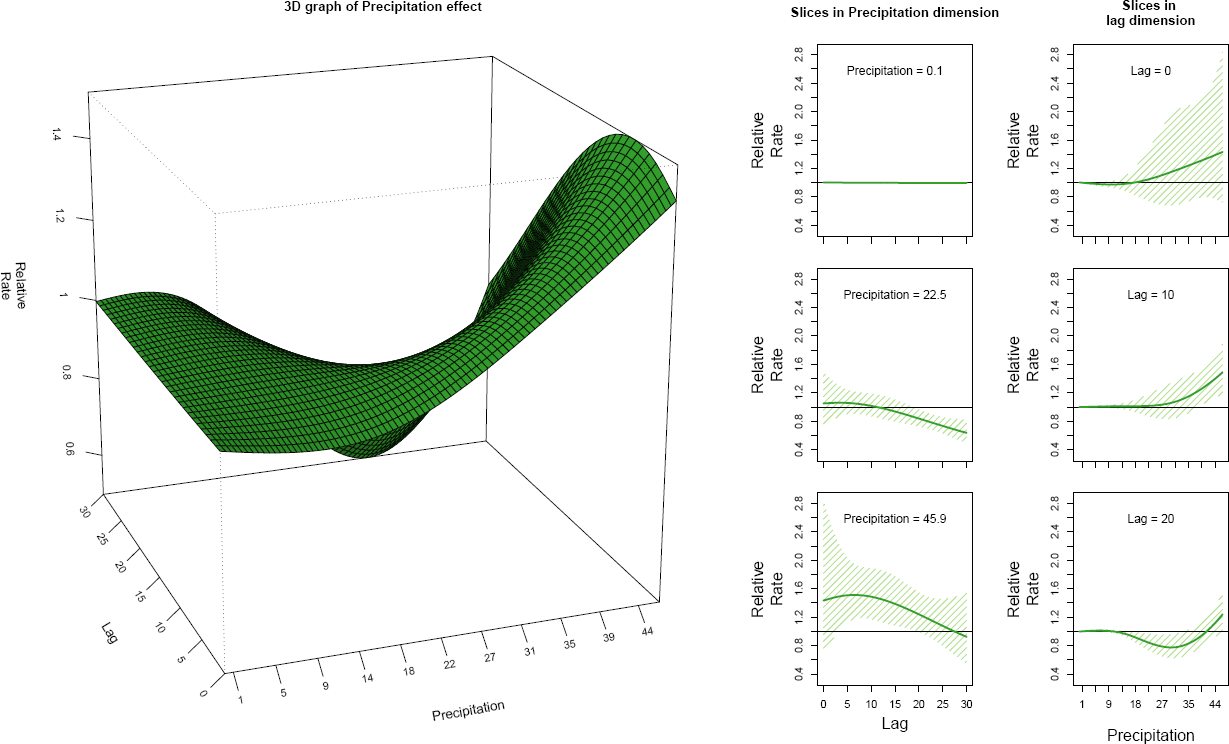
Effect of lagged daily precipitation on mosquito abundance in a univariate model. Panel A: The relationship between precipitation (x-axis), lag up to 30 days (y-axis), and relative rate of number of mosquitoes caught (z-axis) is plotted. Panel B, D, and F plot slices of the surface along the temperature axis with corresponding uncertainty. Panel C, E, and G plot slices of the surface along the lag axis with corresponding uncertainty.

**Figure S8:**
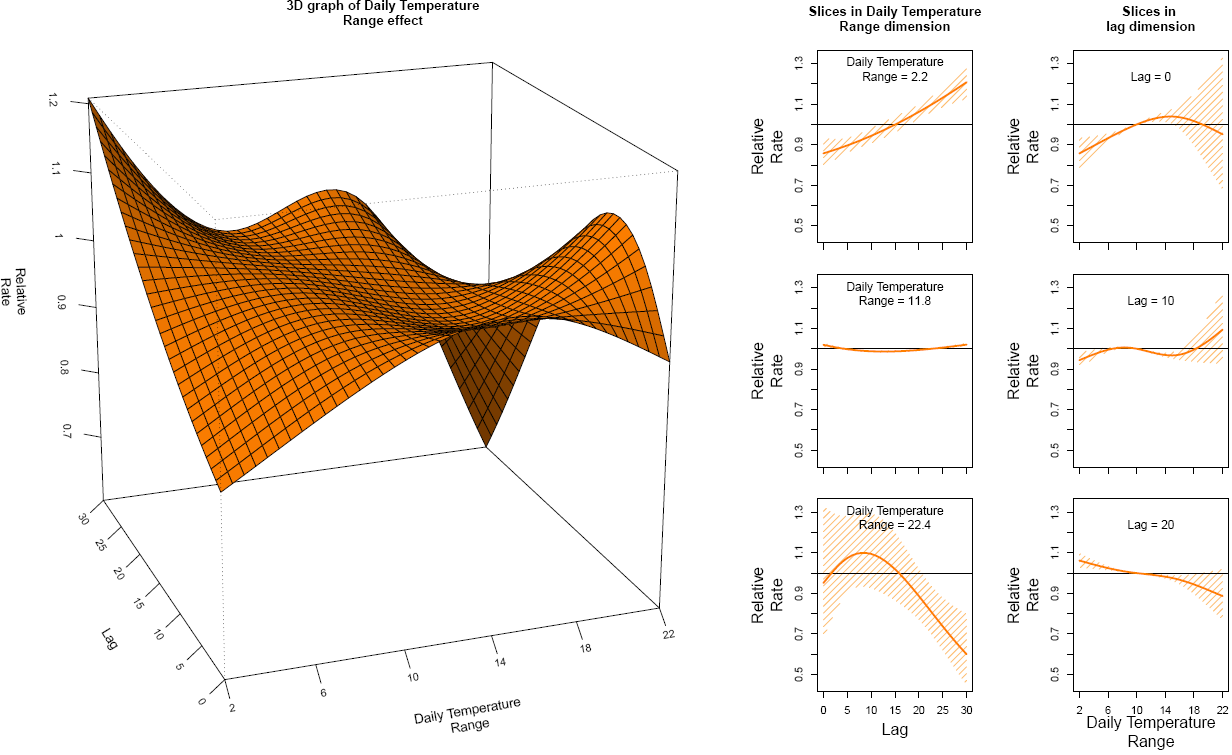
Effect of lagged daily temperature range on mosquito abundance in a univariate model. Panel A: The relationship between daily temperature range (x-axis), lag up to 30 days (y-axis), and relative rate of number of mosquitoes caught (z-axis) is plotted. Panel B, D, and F plot slices of the surface along the temperature axis with corresponding uncertainty. Panel C, E, and G plot slices of the surface along the lag axis with corresponding uncertainty.

**Figure S9:**
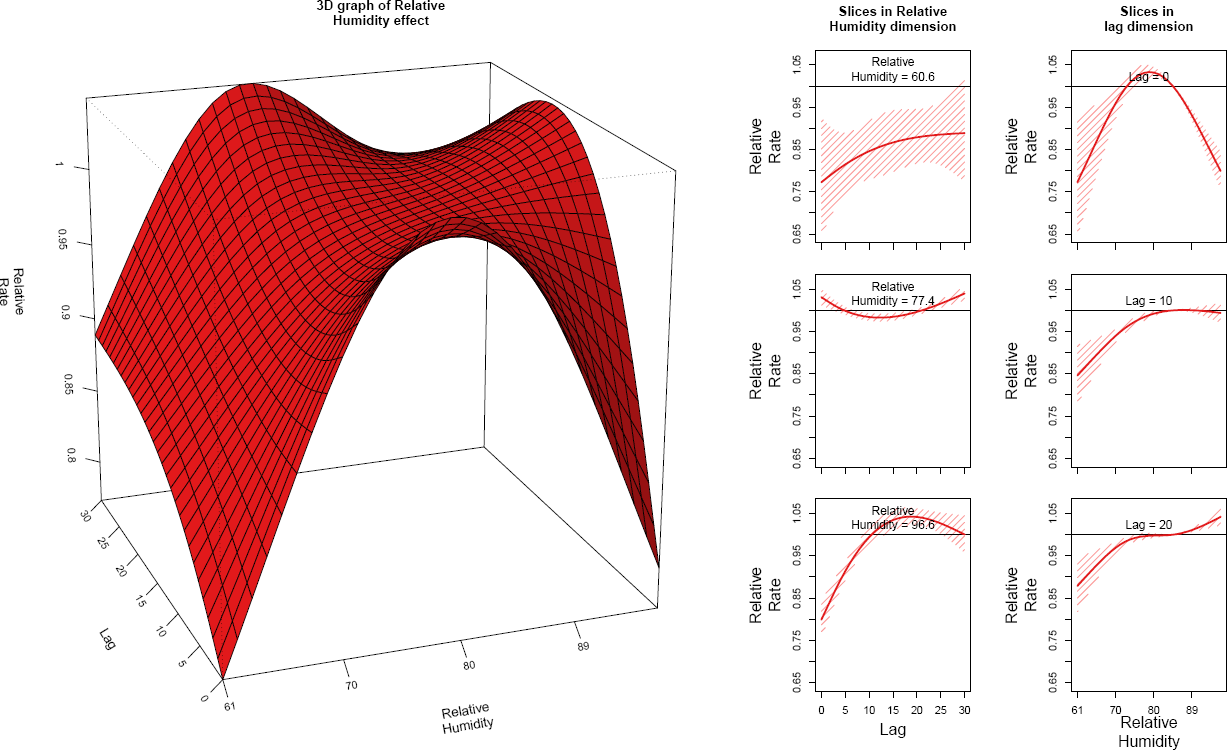
Effect of lagged daily relative humidity on mosquito abundance in a univariate model. Panel A: The relationship between relative humidity (x-axis), lag up to 30 days (y-axis), and relative rate of number of mosquitoes caught (z-axis) is plotted. Panel B, D, and F plot slices of the surface along the temperature axis with corresponding uncertainty. Panel C, E, and G plot slices of the surface along the lag axis with corresponding uncertainty.

**Figure S10:**
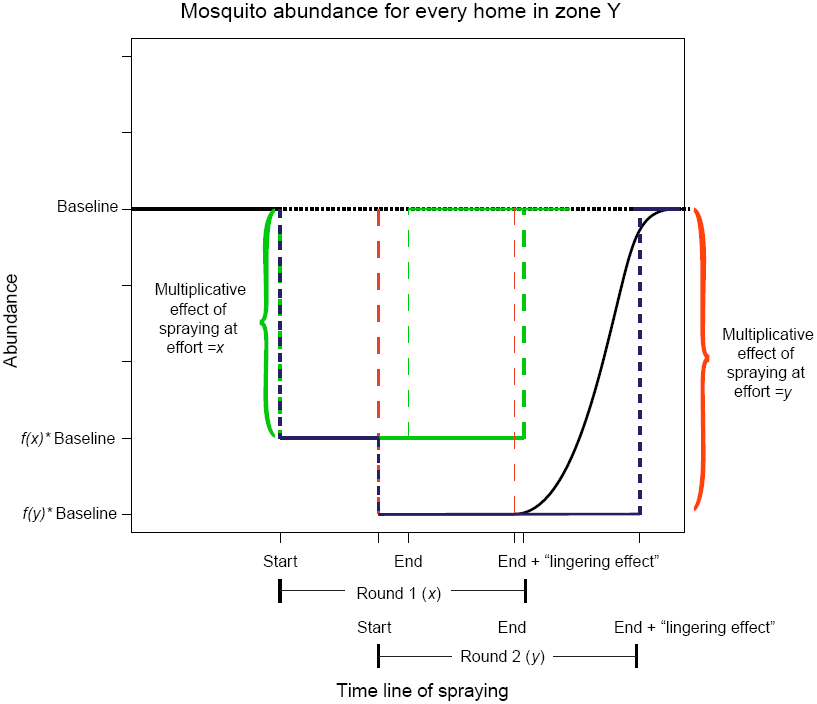
Schematic of the functional form for the impact of intervention coverage on the relative abundance of mosquitoes. The multiplicative effect of the intervention effort reduces mosquitoes from baseline from the beginning of the intervention round through 3 weeks past the end of the intervention round. If two intervention rounds overlap, the effect is assumed to be the larger of each individual impacts and not additive. The SCAM component of the model fits the function *f*(*x*).

**Figure S11:**
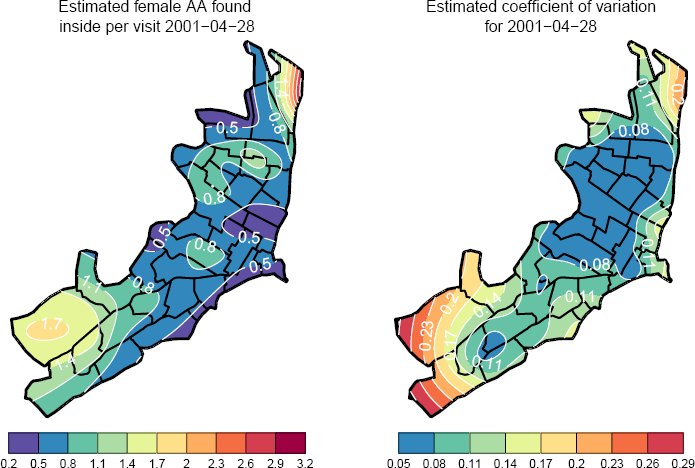
Fitted spatial variation in female *Ae. aegypti* abundance. Panel A: Estimated number of female *Ae. aegypti* that would be caught during a house-hold aspiration held on April 28^th^, 2001 displayed as a contour plot. The choice of this date was arbitrary but necessary to incorporate the climatological covariates. Panel B: Estimated coefficient of variation on number of mosquitoes for April 28^th^, 2001 displayed as a contour plot.

**Figure S12:**
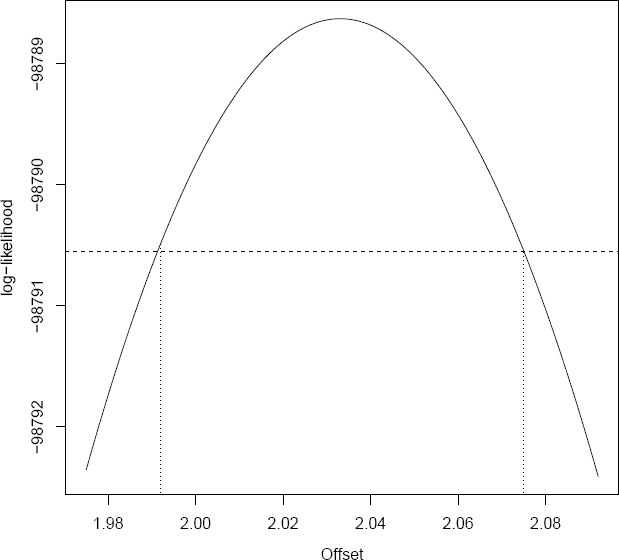
Profile likelihood for the offset. Using the base model, the offset was varied from its maximum likelihood estimate to identify the profile likelihood confidence interval.

**Movie S1: Predicted abundance of mosquitoes through space and time**. Figure 5a is replotted for the first day of each week across the study period. The time-series is representative of the single point in the South-West of Iquitos indicated by an open circle on the 3-D surface.

